# Regional Differences in the Ghrelin-Growth Hormone Secretagogue Receptor Signalling System in Human Heart Disease

**DOI:** 10.1101/2020.07.13.201319

**Authors:** Rebecca Sullivan, Varinder K Randhawa, Tyler Lalonde, Tina Yu, Bob Kiaii, Leonard Luyt, Gerald Wisenberg, Savita Dhanvantari

**Author notes:** **Corresponding Address:** Lawson Health Research Institute, F4-127a, PO Box 5777, Stn B, London, ON, N6A 4V2, Rebecca Sullivan.

## Abstract

The hormone ghrelin and its receptor, the growth hormone secretagogue receptor (GHSR) are expressed in myocardium. GHSR binding activates signalling pathways coupled to cardiomyocyte survival and contractility. These properties have made the ghrelin-GHSR axis a candidate for a biomarker of cardiac function. The dynamics of ghrelin-GHSR are altered significantly in late stages of heart failure and cardiomyopathy, when left ventricular (LV) function is failing. We examined the relationship of GHSR with ghrelin in cardiac tissue from patients with valvular disease with no detectable changes in LV function. Biopsy samples from the LV and left atrium (LA) were obtained from 25 patients with valvular disease (of whom 13 also had coronary artery disease) and preserved LV ejection fraction, and compared control samples obtained via autopsy. Using quantitative confocal fluorescence microscopy, levels of GHSR were determined using a fluorescent peptide analog of ghrelin, Cy5-ghrelin(1-19); ghrelin, the heart failure marker natriuretic peptide type-b (BNP), and contractility marker sarcoplasmic reticulum ATPase pump (SERCA2a) were measured by immunofluorescence. A positive correlation between GHSR and ghrelin was apparent in only diseased tissue. Ghrelin and BNP significantly correlated in the LV and strongly co-localized to the same intracellular compartment in both diseased and control tissue. GHSR, ghrelin and BNP all strongly and significantly correlated with SERCA2a in the LV of diseased tissue only. Our results suggest that the dynamics of the myocardial ghrelin/GHSR axis is altered in cardiovascular disease in the absence of measurable changes in heart function, and may accompany a regional shift in endocrine programming.

## 1. Introduction

Ghrelin is a peptide hormone that has well-known orexigenic effects on the body. Ghrelin is an endogenous 28 amino acid peptide that is a natural ligand of the growth hormone secretagogue receptor (GHSR) 1a, a seven transmembrane G-protein coupled receptor. More recently, ghrelin has been studied for its effects and role in cardiac energetics through activation of myocardial GHSR. Specifically within cardiomyocytes, the binding of ghrelin to GHSR activates downstream signalling pathways that lead to cardioprotective effects, such as maintenance of contractility^1,2^, suppression of inflammation^3,4^, and promotion of growth and survival^5^. Cardiac GHSR activation promotes excitation-contraction coupling, increasing Ca^2+^ flux through the sarcoplasmic reticulum ATPase pump (SERCA2a) and the voltage gated Ca^2+^ channels, which produces a positive inotropic effect and protects against ischemia/reperfusion injury in cardiomyocytes^2^. Ghrelin also regulates inflammation in the heart through the Akt pathway, decreasing expression of the pro-inflammatory markers^6^ IL1-β, TNF-α, and IL-6, and upregulating the TNFα/NFκB pathways^3^. Ghrelin has been shown to protect the heart through the toll like receptor 4 pathway (TLR4)^4^. Apoptosis is decreased upon GHSR activation through extracellular signal-related kinases 1/2 (ERK1/2) and AKT serine kinases^5,6^ and inhibition of caspase-3 and 9^7^. Fibrosis deposition is also minimized in the heart after ghrelin administration^8,9^. Therefore, the ghrelin/GHSR axis within the myocardium plays a critical role in the maintenance of cardiomyocyte function and survival.

The discovery of the myocardial ghrelin-GHSR axis and its role in cardiomyocyte function and health has prompted studies on its dynamics in heart failure and cardiomyopathy. We and others have shown that myocardial GHSR is elevated in end stage heart failure (HF)^10,11^. Additionally, we showed that the cardiac ghrelin-GHSR axis was abnormally up-regulated in end-stage HF compared to endomyocardial biopsies from engrafted hearts^11^, thus suggesting that tissue levels of ghrelin and GSHR could be indicators of the health of transplanted hearts. In contrast, one study has reported decreased GHSR in tissue from patients with end-stage dilated cardiomyopathy, which negatively correlated with LVEF^12^. We^13^ and others^14^ have shown that tissue GHSR also decreased in mouse models of diabetic cardiomyopathy, suggesting that the dynamics of the myocardial ghrelin-GHSR axis may differ in earlier stages of myocardial decompensation as compared with end-stage heart failure.

Some potential causes of heart failure include coronary artery disease (an accumulation of cholesterol (plaques) in the vessel walls) or aortic stenosis (progressive restriction of the opening of the aortic valve)^15^, and there has been some clinical investigation into correlations with serum ghrelin levels or the effects of ghrelin administration on heart function in these conditions. In humans with coronary artery disease with diabetes-associated coronary atherosclerosis, serum levels of ghrelin decreased significantly compared to healthy controls^16,17^. Another study also showed an inverse relationship between serum ghrelin and coronary artery disease (CAD) detected by angiography that was independent of other cardiovascular risk factors^16^, suggesting that a reduction in ghrelin-GHSR signalling may be correlated with progressively more severe CAD. Ghrelin administration in rats with aortic stenosis reduced calcification buildup in the myocardium in a dose-dependent manner^18^, and attenuated aortic calcium deposition both *in vivo* and *in vitro^19^*. Therefore, investigating the dynamics of the ghrelin-GHSR axis and signaling within cardiomyocytes may provide further insight into the potential use of ghrelin as a therapy for myocardial dysfunction (heart failure).

In this study, towards the goal of determining the potential of ghrelin-GHSR as a biomarker of heart disease, we examined their levels in a cohort of patients who underwent elective cardiac valve replacement surgery and compared their expression levels to control heart tissues. We also determined the correlation of ghrelin-GHSR with an array of biomarkers including natriuretic peptide type-B (BNP), sarcoplasmic reticulum ATPase (SERCA2a), and toll-like receptor 4 (TLR4). With these data, we hope to obtain further insights into the distinct molecular signalling patterns associated with the progression of heart disease.

## 2. Materials and methods

### 2.1 Patient Cohort

Tissue samples were harvested from 25 patients who underwent aortic valve replacement surgery for aortic stenosis at the London Health Sciences Center (LHSC) between 2013 and 2014. Twenty-four of these patients had aortic stenosis and one had mitral valve impairment. Of these 25 patients, 11 concurrently had coronary artery disease and received coronary artery bypass grafts. Sixteen patients were male and nine were female, with an overall average age of 66 years. Twenty-three patients had normal ejection fractions (>45%) prior to surgery, and 2 patients had reduced ejection fractions (35-40%). The protocol for obtaining tissue samples was approved by Western University’s Health Sciences Research Ethics Board. Tissue samples, roughly 0.2cm to 0.5cm in length, were collected from the left atrial (LA) and left ventricular (LV) myocardium from each patient. The left atrial samples were obtained from the posterior left atrial wall just on the left atrial side of the inter-atrial septum (close to the right atrium but technically the left atrium). The left ventricular samples were taken from the posterior left ventricular wall, just below the mitral valve. Patient demographics, cardiac function (LVEF), and medications are shown in Tables 1 and 2. All patient samples and patient data were kept anonymous and all biomarker analyses were completed prior to receiving clinical data.

**Table 1.** Cardiac Surgery Patient Demographics

| Patient Condition | Number of Males with Condition (n of 16) | Number of Females with Condition (n of 9) |
| --- | --- | --- |
| Age range (years) | 49 – 84 | 45 – 77 |
| Mean age (years) | 69 | 60 |
| Coronary artery disease | 9 | 2 |
| Moderate – Severe Aortic Stenosis | 14 | 8 |
| Hypertension | 8 | 2 |
| High pulmonary artery pressure | 10 | 7 |
| Diabetes | 2 | 1 |
| LVEF <45% | 2 | 0 |
| LVEF >45% | 14 | 9 |

**Table 2.** Patient Medications Post-Heart Surgery.

| Patient Medication | Number of Males with Condition (n of 16) | Number of Females with Condition (n of 9) |
| --- | --- | --- |
| ACE Inhibitor | 1 | 2 |
| Anti-arrhythmic | 1 | 2 |
| Angiotensin receptor blocker (ARB) | 6 | 1 |
| Coumadin | 4 | 2 |
| Beta blocker | 5 | 5 |
| Calcium channel blocker | 2 | 3 |
| Diuretics | 8 | 3 |
| Statins | 11 | 3 |

In order to provide a reference non-diseased control, paraffin-embedded and sectioned samples of cardiac tissue (n=10) were obtained from the tissue archives of the Department of Pathology, London Health Sciences Centre. These tissue samples had been obtained from the LV of patients who had died from non-cardiac causes at the time of post-mortem, 24-48 h after the patient was pronounced dead. Of these 10 patients, 5 were male and 5 were female. The average age of all ten patients was approximately 59 years. Analysis of these tissues was compared to the analysis derived from the surgery patients for overall marker fluorescence intensity and differences in biomarker relationships.

### 2.2 Quantitative Fluorescence Microscopy

Samples were fixed, frozen and embedded in optimal cutting temperature compound (OCT), and subsequently sectioned at 6-7 μm thickness, as previously described^20,21^. Immunohistochemistry using primary and fluorophore-conjugated secondary antibodies was conducted as previously described^20,21^. In brief, tissue sections were incubated with blocking buffer in 10% serum for 30 minutes at room temperature followed by incubation with primary polyclonal or monoclonal antibodies (Table 3) for 1h at room temperature in a humidified chamber. These antibodies were used to identify ghrelin (1:100) [Santa Cruz Biotechnology], BNP (1:1000) [Abcam], SERCA2a (1:300) [Abcam], and TLR4 (1:250) [Abcam]. Samples were rinsed twice in phosphate buffered solution (PBS) and incubated for 2h at room temperature with secondary antibodies (1:500) (Table 3). To detect GHSR, we used the far-red ghrelin peptide analog, [Dpr^3^(n-octanoyl),Lys^19^(sulfo-Cy5)]ghrelin(1-19), referred to as Cy5-ghrelin(1-19), as we have previously done to quantify GHSR *in situ^12^*. This analog binds with high specificity to GHSR in mouse and human cardiac tissue samples^11,13^. After incubation with secondary antibodies, Cy5-ghrelin(1-19) was added to tissue sections for 30 min. Sections were washed twice with PBS, incubated 8 min with DAPI nuclear stain (1:1000), and mounted with ProLong Diamond antifade (Life Technologies) to prevent the tissues from photobleaching. N numbers for each biochemical marker were different due to the limited amount of tissue available at the time of staining.

**Table 3.** Antibody Table – Information on Antibodies Used

| Antigen | Catalog Number | Dilution | Host |
| --- | --- | --- | --- |
| Ghrelin | sc-10359 | 1:100 | Goat |
| BNP | ab19645 | 1:1000 | Rabbit |
| SERCA2a | ab3265 | 1:300 | Rabbit |
| TLR4 | ab22048 | 1:250 | Mouse |
| DAPI | 62247 | 1:1000 | - |
| Alexa Fluor 488 | A11055 | 1:1000 | Donkey anti-goat |
| Alexa Fluor 594 | A21207 | 1:1000 | Donkey ant-rabbit |
| Alexa Fluor 488 | A21206 | 1:1000 | Donkey anti-rabbit |
| Alexa Fluor 594 | A21203 | 1:1000 | Donkey anti-mouse |

For control cardiac tissue, all sections were deparaffinized with decreasing concentrations of alcohol solutions (95%-70%). Samples were then stained as previously stated for the following biomarkers: ghrelin (1:100) [Santa Cruz Biotechnology], BNP (1:1000) [Abcam], SERCA2a (1:300) [Abcam], and TLR4 (1:250) [Abcam] and GHSR using Cy5-ghrelin(1-19) (1:100).

### 2.3 Background subtraction for fluorescence microscopy imaging

To ensure specificity of the fluorescent signal for GHSR, background subtraction was performed, as GHSR has a diffuse staining pattern. To calculate the background signal, tissue sections were first incubated with 100μM hexarelin, a GHSR ligand^22^ known to competitively displace Cy5-ghrelin(1-19)^21^, for 1 hour at room temperature prior to incubation with 10μM Cy5-ghrelin(1-19) as described above. Tissues were then stained for DAPI (1:1000), washed and mounted with ProLong Diamond antifade as described above. For control tissue samples, background fluorescence intensity values were calculated in 5 separate control samples, and the average value of these intensities was subtracted from the total fluorescence intensities of GHSR in each of the 10 original tissue samples. For diseased tissue samples, background values were calculated from 5 fields of view in one slide from each patient, and the average value of this slide was calculated as the background fluorescence. These values were subtracted from the total fluorescence intensities of non-blocked tissue from that patient to obtain specific Cy5-ghrelin(1-19) signal in the diseased tissue.

### 2.4 Image Acquisition and Analysis

High resolution images of both the diseased and control tissues were captured with a Nikon A1R Confocal Microscope at 60x magnification using an oil immersion lens. Five random fields of view were acquired for each of 2 tissue sections per patient sample with the exposure time, gain and LUT set the same for all tissue sections. Therefore, each patient sample (reported as data points in all graphs) reflects an average of 2 technical replicates.

Images of GHSR, ghrelin, BNP, SERCA2a, and TLR4 were analyzed with FIJI 1.49v, a distribution of ImageJ software (National Institutes of Health, Bethesda). Algorithms built into ImageJ were used to quantify different staining patterns in the tissue. Punctate staining patterns were quantified using the RenyiEntropy algorithm, an entropy-based approach that distinguishes the positive punctate signals within the cells from the background. The integrated density represents the mean intensity of the positive signal above a certain threshold in scaled units divided by the area in pixels. The integrated density threshold was set such that the entropies of distributions above and below the set threshold are maximized. Therefore, this algorithm only captured the high intensity punctate staining patterns and did not calculate any background staining of these images. A representation of what the algorithm identified as positive versus background is shown in Supplemental Figure S1.

For diffuse staining patterns, the Li algorithm was used to quantify the specific fluorescence signal after background subtraction as described above. All images for diffuse staining were quantified using the built-in ImageJ Li algorithm with the integrated density (stated above) determined for each tissue.

### 2.5 Fibrosis Imaging

To assess fibrosis, heart tissue sections were stained with Masson’s Trichrome stain by the core Pathology Laboratory at LHSC. Sections of both the diseased and control tissue were acquired using bright field microscopy at 10X and 20X magnifications with a Zeiss Axioskop EL-Einsatz microscope and Northern Eclipse software as described previously^11,13^. Images were acquired for the entire tissue section and quantification was performed on the entire tissue.

Fibrosis was analyzed using an online script in the program ImageJ Fiji which quantified the percentage of fibrotic tissue in each sample by distinguishing fibrotic tissue (blue) from non-fibrotic tissue (red), as previously described^23^. Briefly, this script divides the RBG image into each of its channels which thresholds the image for positive pixels in either the red or blue channels. All tissue sections were set at the same threshold for both the fibrotic and non-fibrotic tissue. The fibrosis was then quantified based on the fibrotic tissue compared to the total area of tissue to determine the average percentage of fibrosis in each of the diseased or control tissue samples.

### 2.5 Data Analysis

Data from biomarkers (GHSR, ghrelin, BNP, SERCA2a, and TLR4) were disaggregated by cardiac region using a post-hoc analysis. Statistical analyses were performed using GraphPad Prism version 8. Unpaired student *t*-tests were used to compare overall biomarker expression differences between the diseased and control tissue. Within the diseased tissue cohort, a one-way ANOVA with analysis of variance using Tukey *post-hoc* test was used to determine regional differences. Linear regression was used to determine correlations between the following biomarkers: GHSR, ghrelin, BNP, SERCA2a and TLR4. Linear regression analysis was used to compare biomarker fluorescence intensities based on the following categories: entire patient cohort and regional differences (left atrium and left ventricle). Pearson correlation was used to determine colocalization between ghrelin and BNP in both diseased (LA and LV) and control tissue. All statistical analyses were completed with significance set at p<0.05.

## 3. Results

### 3.1 Cardiac surgery patient cohort

Of the twenty-five patients who underwent cardiac surgery, 16 were male and 9 were female with mean ages of 69 and 60 respectively (Table 1). The patients had a range of moderate to severe aortic stenosis while 11 also had coronary artery disease. In the patients with coronary artery disease, there were no correlations between any markers evaluated in this study (GHSR and ghrelin – r=0.3942 p=0.1826; GHSR and BNP – r=0.0596 p=0.8467; ghrelin and BNP – r=0.1769 p=0.8850; SERCA2a and GHSR – r=0.4132 p=0.2354; SERCA2a and ghrelin – r=0.5175 p=0.1256; SERCA2a and BNP – r=0.4315 p=0.2131). Sixteen patients had elevated estimated pulmonary artery pressures, based on their pre-operative echocardiograms and only two patients had a reduced ejection fraction prior to surgery (30-40%). After cardiac surgery, 10 patients were on beta blockers, 11 were on diuretics and 14 were on statins (Table 2).

### 3.2 Correlations Between GHSR and Ghrelin in Diseased Cardiac Tissue

Fluorescence intensities of both GHSR and ghrelin were variable in any given cardiac tissue sample in both the diseased and control cohort (Fig 1A & C). Overall GHSR expression was not significantly different between the diseased and control groups (Fig 1B) while tissue ghrelin was significantly lower (p<0.0001) in the diseased cohort in both the LA and the LV (Fig 2 D). Linear regression analysis of GHSR vs. ghrelin showed a positive relationship in the overall cardiac surgery cohort (r = 0.3995; p=0.0318) (Fig 2 C). Regionally, this correlation persisted only in the left atrium (r = 0.4859; p=0.0299) and not in the left ventricle (Fig 3D). There was no significant correlation between GHSR and ghrelin in the control tissues (r = 0.1105; p=0.3480) (Fig 2B).

**Figure 1.**
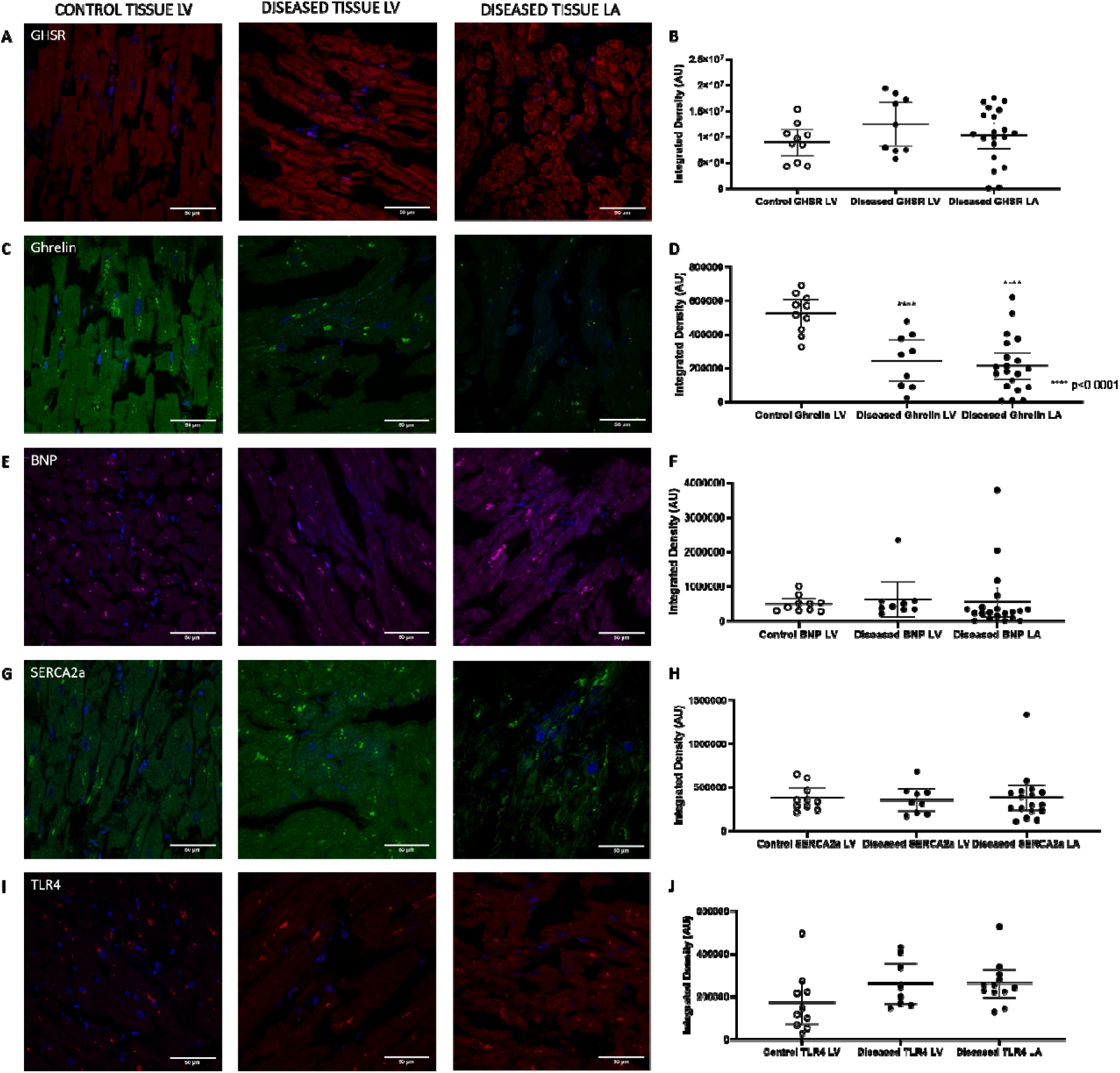
Representative confocal fluorescence images of all biomarkers (Cy5-ghrelin(1-19) – red, ghrelin – green, BNP – magenta, SERCA2a – green, TLR4 – red) in both the control tissue LV (left column) and diseased tissue LV (middle column) and LVA (right column). All images indicate DAPI nuclear stain in blue. Graphs represent quantification of fluorescent images as means ± 95% CI of integrated densities for each biomarker with each dot representing one patient sample. There were no significant differences in fluorescence intensities of (B) Cy5-ghrelin(1-19) [control LV n=10, LV n=9, LA=20], (F) BNP [control LV n=10, LV n=9, LA=20], (H) SERCA2a [control LV n=10, LV n=9, LA=17] or (J) TLR4 [control LV n=10, LV n=8, LA=12]. There was a significant decrease in the fluorescence intensities detecting (D) ghrelin (p<0.0001) [control LV n=10, LV n=9, LA=20] in the diseased cohort compared to the control tissues.

**Figure 2.**
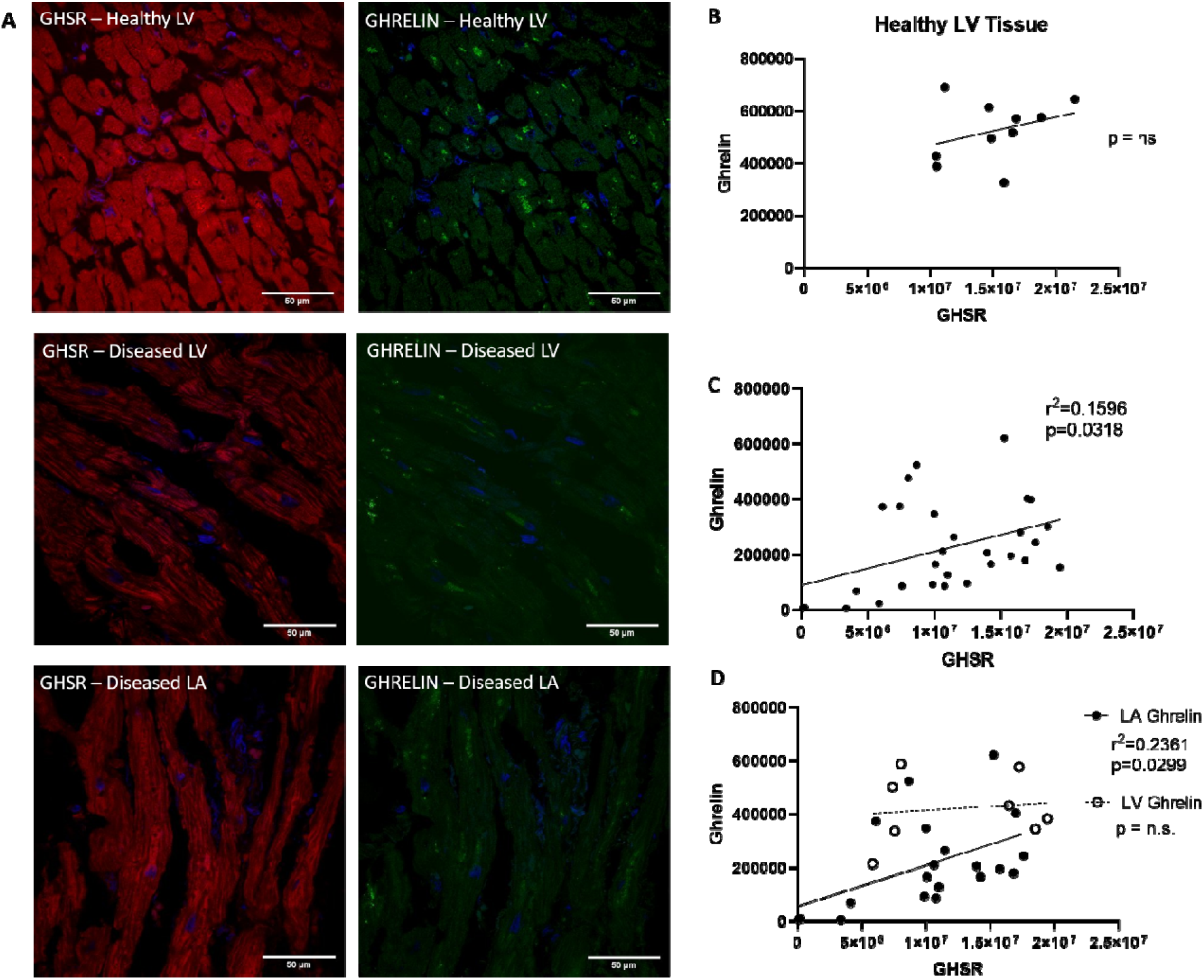
GHSR and ghrelin fluorescence intensity in human cardiac tissue. (A) Representative confocal images of Cy5-ghrelin(1-19) (red), and ghrelin (green) in diseased control LV (top), diseased LV (middle) and diseased LA (bottom). DAPI (blue) indicates nuclear localization in cardiomyocytes. Graphs indicate quantification of integrated densities using linear regression between Cy5-ghrelin(1-19) and ghrelin. In control tissue, there was no correlation between Cy5-ghrelin(1-19) and ghrelin (p=ns) [n=10] (B). Linear regression between Cy5-ghrelin(1-19) and ghrelin (r = 0.3995; p=0.0318) in the entire patient cohort [n=29] where each dot represents one patient sample (C). Regional analysis showed a significant linear regression of Cy5-ghrelin(1-19) and ghrelin in the LA (r = 0.4859; p=0.0299) [n=20] but not the LV (p=ns) [n=9] (D). LA, Left Atrium; LV, Left Ventricle.

**Figure 3.**
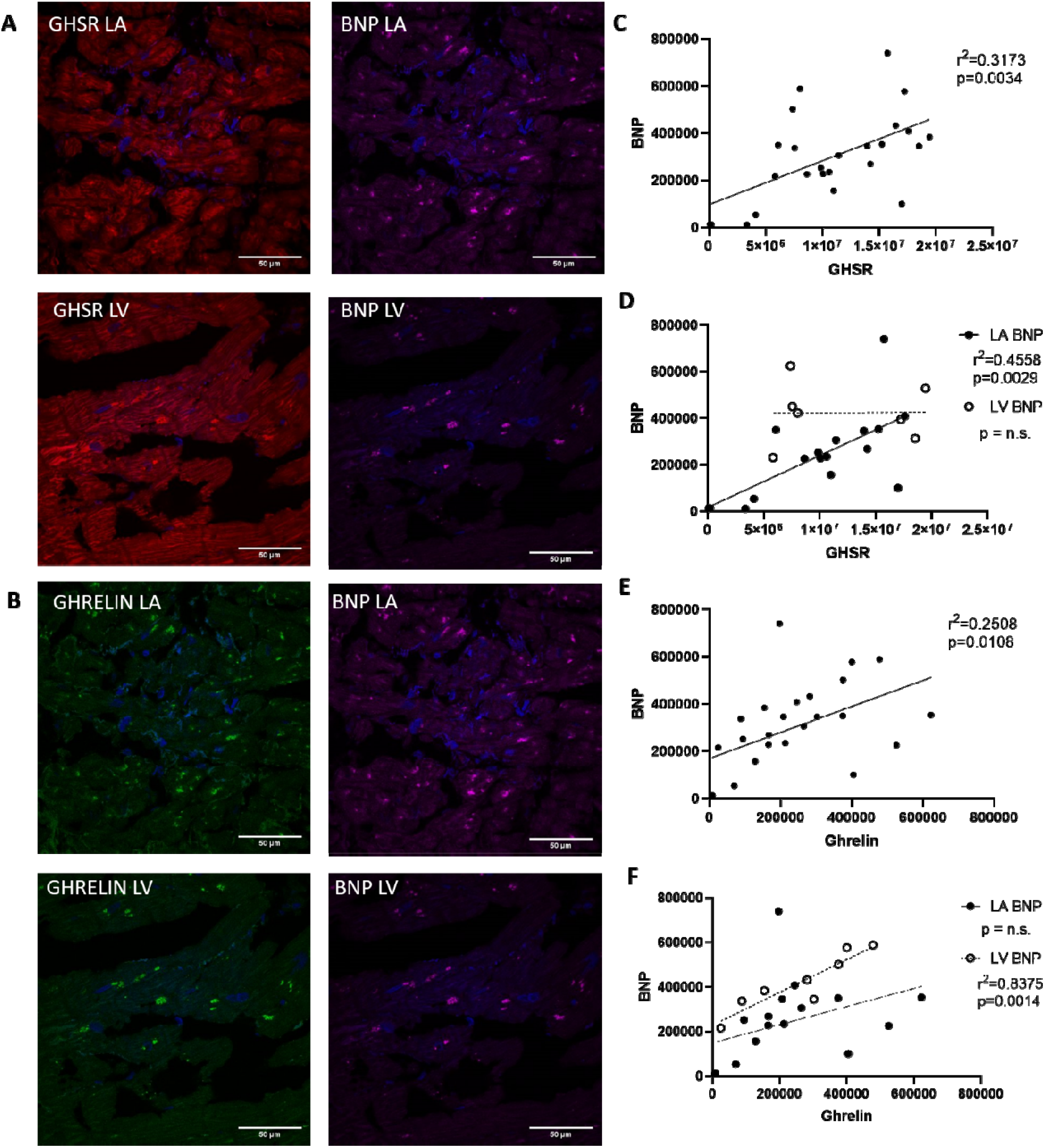
Representative fluorescent confocal images of Cy5-ghrelin(1-19) (red), ghrelin (green) and BNP (magenta) in human cardiac tissue in the LA and LV (A & B). Nuclei were visualized with DAPI (blue). Graphs show linear regression analysis of quantified integrated densities with each dot representing an individual patient sample. There was significant linear regression of Cy5-ghrelin(1-19) vs BNP (r = 0.5633; p=0.0034) in the entire patient cohort [n=25] (C). When data were disaggregated by region, a significant linear regression was found in the LA (r = 0.6751; p=0.0029) [n=17] but not the LV (p=ns) [n=8] (D). Ghrelin and BNP were correlated in the entire patient cohort (r = 0.5008; p=0.0108) [n=25] (E). Linear regression analysis between ghrelin and BNP indicate correlations in the LV (r = 0.9152; p=0.0014) [n=17] but not the LA (p=ns) [n=9] (F). LA, Left Atrium; LV, Left Ventricle.

### 3.3 Relationship of BNP to GHSR and Ghrelin in Diseased Cardiac Tissue

Fluorescence images indicated the localization of BNP within the diseased and control heart tissue (Figure 1E). Analysis of these images indicated no significant difference in immunofluorescence (p=ns) in the disease cohort compared to the control tissues (Fig 1F). Linear regression analysis indicated a strong and significant positive relationship between BNP and GHSR (r = 0.5633; p=0.0034); this relationship was strong within the LA (r = 0.6751; p=0.0029) but not the LV (r = 0.0127; p=0.9784) (Fig 3 C – D). There was also a significant positive relationship between BNP and ghrelin in the entire cohort (r = 0.5008; p=0.0108). This relationship, however, differed by region: LV (r = 0.9152; p=0.0014), LA (r = 0.394 p=0.1177) (Fig 3 E – F). Again, no relationship was seen between BNP and GHSR or ghrelin in the control cardiac tissue (r = 0.2465; p=0.4923 and r = 0.1236; p=0.7337 respectively).

### 3.4 Intracellular Colocalization of Ghrelin and BNP in cardiomyocytes

Ghrelin and BNP appeared to be geographically localized to the same intracellular compartment within cardiomyocytes. In the control tissue, ghrelin and BNP appeared as larger ordered punctate areas mainly surrounding the nucleus (Fig 4A) while in the diseased tissue these colocalized areas of ghrelin and BNP were smaller and scattered more diffusely throughout the cell, and are not focused primarily around the nucleus (Fig 4B – C). The Pearson Correlation Coefficient was strong for both diseased LA (PCC=0.76) and LV (PCC=0.83) and control (PCC=0.86) while there was a significantly higher correlation in the control tissues compared to the LA (p=0.00169) diseased tissue but not LV (p=0.2923) (Fig 4D).

**Figure 4.**
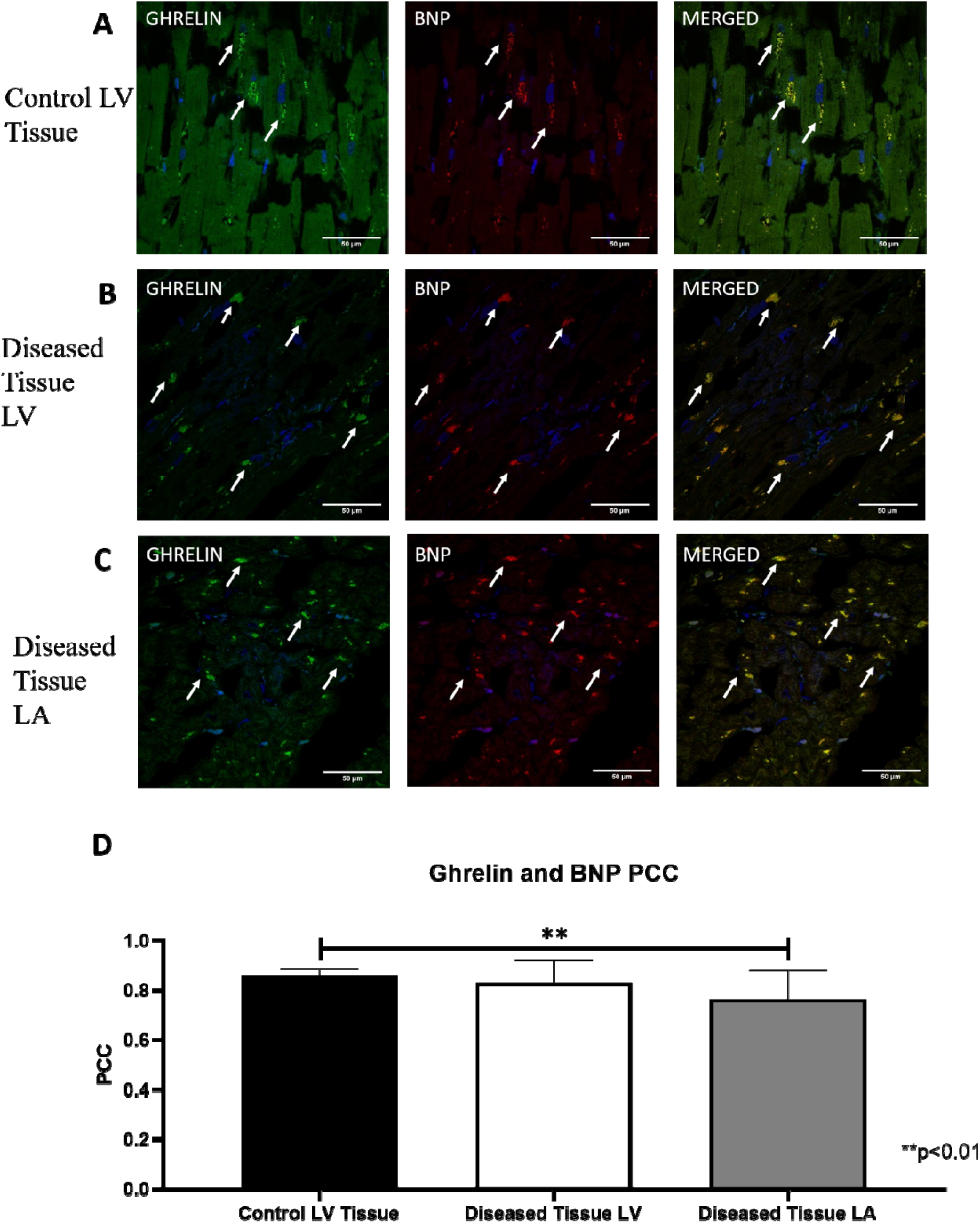
Intracellular colocalization of ghrelin and BNP in control LV (A) and diseased tissue LV (B) and LA (C). (A-C) Representative confocal fluorescent images of ghrelin (green) and BNP (red) in healthy and diseased tissue shows correlation of ghrelin and BNP in the merged images. DAPI (blue) indicates nuclear staining. White arrows indicate areas of punctate staining and colocalization in the merged images. (D) Pearson Coefficient Correlation (PCC) values indicate a stronger correlation in the healthy tissue [n=10] compared to the diseased tissue in the left atrium (p=0.0040) [n=29] but not the left ventricle (p=ns) [n=29]. Values are means ± SD.

### 3.5 The Relationship of the Cardiac Contractility Biomarker SERCA2a to Ghrelin and GHSR in the Diseased Heart

Quantitative fluorescence microscopy was used to measure SERCA2a in the control and diseased cohorts with representative images shown in Fig 1G. No differences in overall expression were seen between the diseased and control cohorts (Fig 1H). Linear regression analysis showed that SERCA2a had a positive relationship to GHSR, ghrelin and BNP (Fig 5C – E) but only in the LV (r = 0.7893, p=0.0348; r = 0.7315, p=0.0392; r = 0.8401, p=0.0180 respectively) and not the LA. Quantification of overall expression of toll-like receptor 4 (TLR4) showed no differences between the diseased and control cohorts (Fig 1J). No relationships are present in the control group between SERCA2a and any other biomarker [GHSR (r = 0.0936; p=0.7970), ghrelin (r = 0.0089; p=0.9385), BNP (r = 0.2492; p=0.4875)].

**Figure 5.**
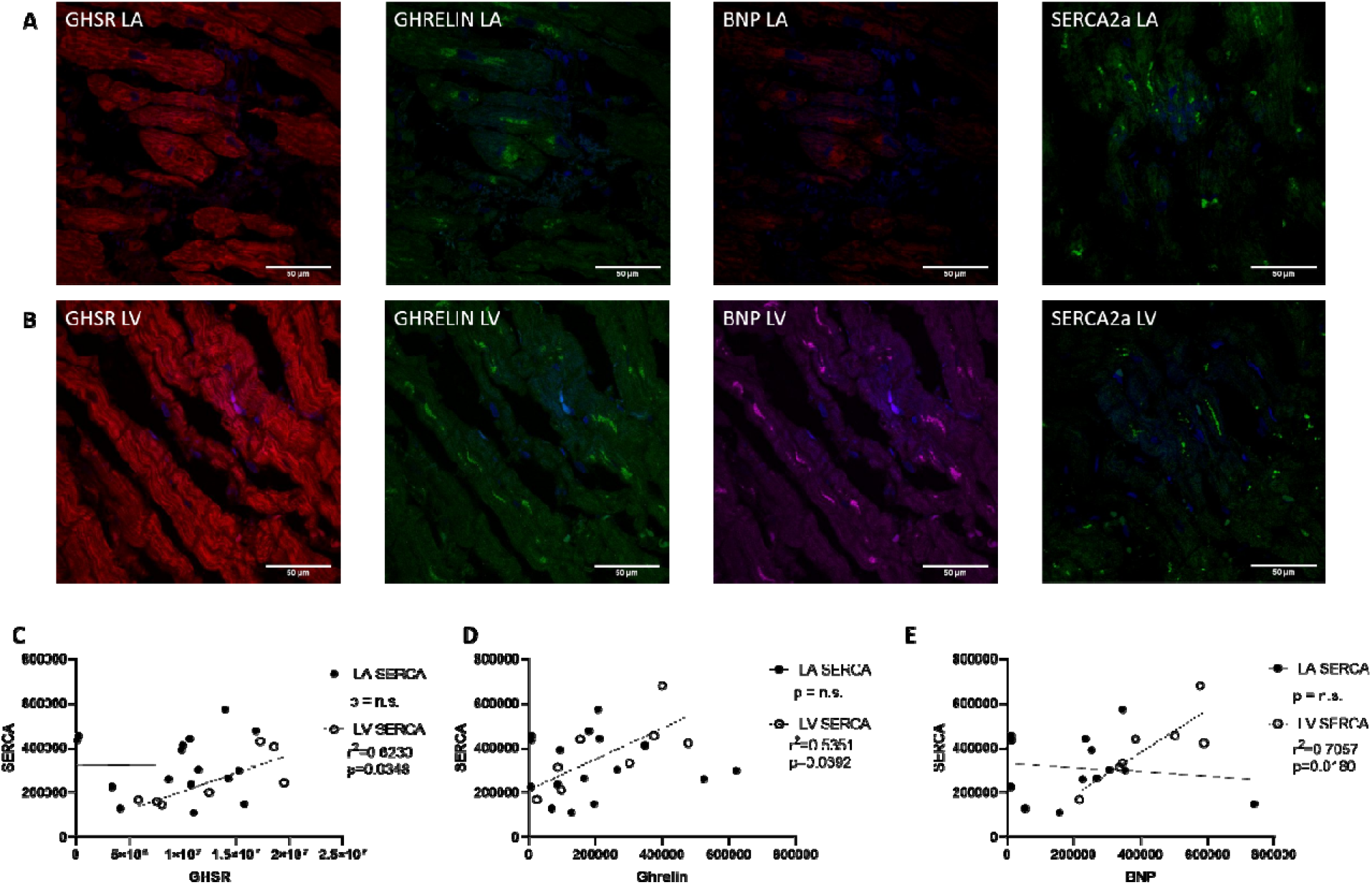
(A) Representative fluorescent confocal images of Cy5-ghrelin(1-19) (red), ghrelin (green), BNP (magenta), and SERCA2a (green) in cardiac tissue of the LA (A in top panels) and the LV (B middle panels). DAPI (blue) nuclear stain in all images. Graphs indicate fluorescence intensities represented by integrated densities where each dot represents one patient sample. A significant linear regression is present in only the LV between SERCA2a and (C) Cy5-ghrelin(1-19) (r = 0.7893; p=0.0348) [LA n=15, LV n=8], (D) ghrelin (r = 0.7315; p=0.0392) [LA n=15, LV n=8], and (E) BNP (r = 0.8401; p=0.0180) [LA n=14, LV n=7]. LA, Left Atrium; LV, Left Ventricle.

### 3.5 Cardiac Fibrosis

Masson’s trichrome staining was used to determine the extent of fibrosis in all cardiac tissue samples. There were highly variable amounts of fibrotic deposition (blue) in samples of diseased tissue compared to the non-fibrotic tissue (red) in any given cardiac surgery patient sample (Fig 6A – C) but not in the control tissue. Quantification of the amount of fibrotic tissue showed a significantly higher amount of collagen deposition in diseased tissues of the LA (p=0.0034) but not of the LV (p=0.0509) compared to the control tissues (Fig 6D).

**Figure 6.**
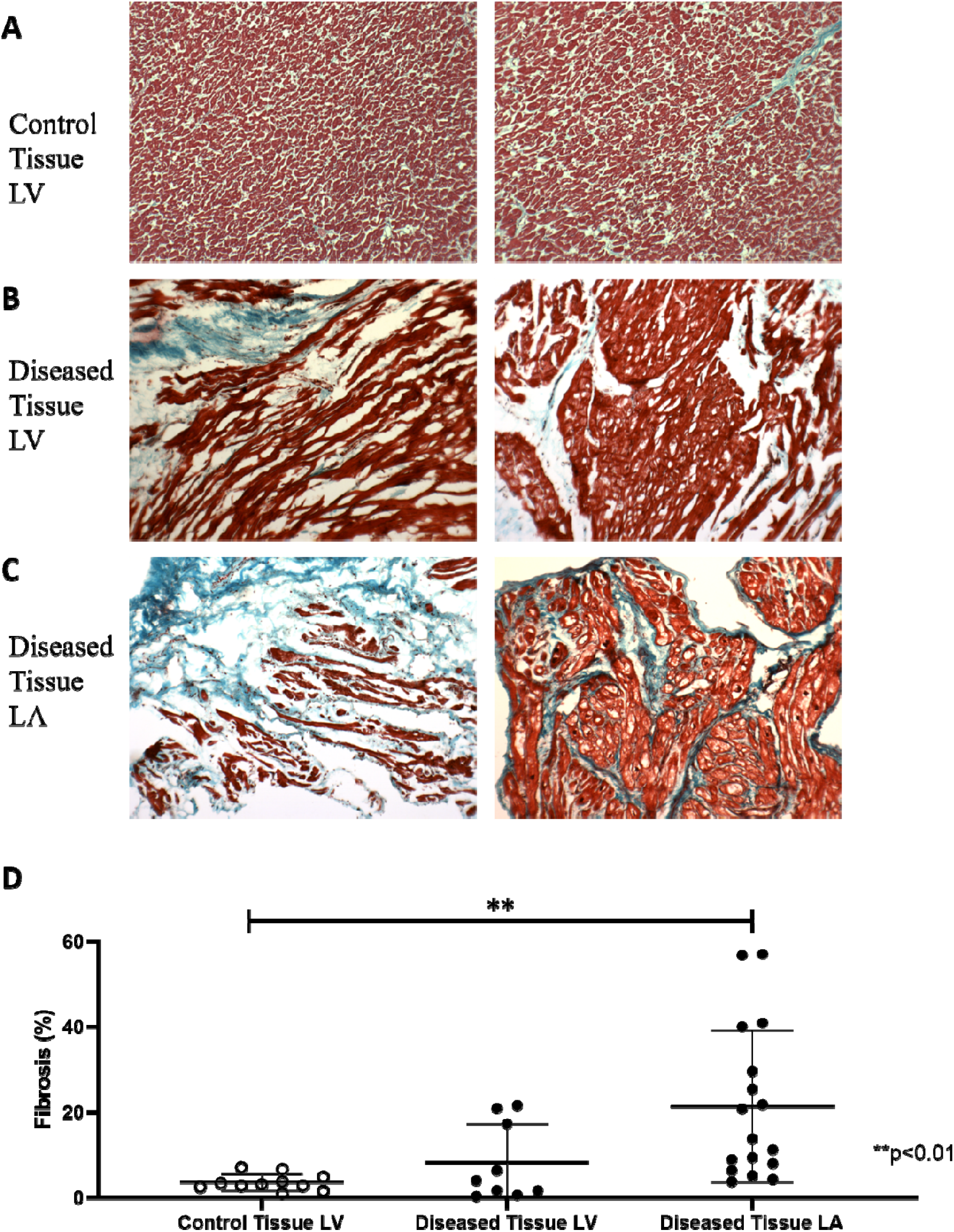
Fibrosis deposition in healthy and diseased tissue. Representative images of fibrotic (blue) and non-fibrotic tissue (red) in control tissue LV (A) and diseased tissues of the LV (B) and the LA (C) show variability in fibrotic deposition in the diseased tissue while not in the control tissue. (D) Graph shows mean ± 95% CI of percentage fibrotic tissue with significantly less fibrotic deposition in the control tissue [n=10] compared to the left atrium (p=0.0034) [n=17] but not the left ventricle (p=ns) [n=11].

## 4. Discussion

In this study, we used our fluorescent ghrelin analog, Cy5-ghrelin(1-19), to examine the expression of myocardial GHSR in relation to that of ghrelin and other known markers of downstream signaling pathways in tissue obtained from patients with valvular heart/coronary artery disease but without reduced LVEF. We also measured these markers in control, nondiseased, cardiac tissue. In patients with valvular heart disease (±coronary artery disease), there were positive correlations between GHSR and ghrelin which were regionally constrained to the left atrium. Similarly, there were regionally divergent correlations between GHSR and BNP (the gold standard marker of HF) and ghrelin and BNP. The positive correlations with the contractility marker SERCA2a were specific to the left ventricle. In contrast, no correlations between GHSR and ghrelin, or any other biochemical marker were observed in the control tissue. Additionally, we found that both ghrelin and BNP localized to the same intracellular compartment within cardiomyocytes, and this colocalization was slightly disrupted in the left atrium in the heart disease (HD) cohort. Therefore, there was an emergence of region-specific patterns in the myocardial ghrelin-GHSR signalling axis in patients with valvular disease despite the absence of measurable changes in heart function.

We have previously shown that GHSR and ghrelin are positively correlated in tissue samples from patients with end-stage heart failure and are correlated negatively with LVEF^11^. Results from other studies also indicate changes in the dynamics of the ghrelin/GHSR axis in end-stage heart failure^10^ and dilated cardiomyopathy^12^, suggesting this cardioprotective ghrelin-GHSR system is abnormally up-regulated when there is injury or stress to the heart^11^. Our present results also show a positive correlation between ghrelin and GHSR in patients with heart disease, but this time in the absence of a decrease in LVEF. As this correlation was not observed in tissue samples from the control group, we suggest that changes in the myocardial ghrelin-GHSR axis occur early in the progression of heart disease, (in this study, in hearts subject to increased LV systolic pressures secondary to aortic stenosis) prior to measurable changes in heart function, as defined by the global LV ejection fraction.

Post-hoc disaggregation of data by region showed that the positive correlation between ghrelin and GHSR was maintained in the left atrium (LA) but not the left ventricle (LV). Interpretation of these data is very limited as we did not collect data on the LA pressures which are commonly elevated in valvular diseases due to stretching of the atrial wall. We did, however, observe a very slight but significant negative correlation between GHSR in the left atrium and LV diastolic pressure (r = 0.6887, p = 0.0402). Left ventricular diastolic pressure is elevated in valve disease which is associated with increased LA pressure^24^. In one study by Schwarz *et al*., in patients with aortic valve disease, there was an elevation in end-diastolic left ventricular pressure and mean left atrial pressure along with significant correlations with myocardial cell diameter^25^. Therefore, in our study of patients with aortic stenosis, it is expected that with the increased LV diastolic pressure, left atrial pressures are also increased. The increased LA pressures may be related to the strong correlation between ghrelin and GHSR. More robust data will be required to establish a concrete relationship between alterations in the ghrelin-GHSR axis and onset of LA dysfunction.

In addition to the myocardium, ghrelin and GHSR are also present in the endothelial cells of the aorta, coronary arteries, and pulmonary arteries and veins^26^. Ghrelin is known to have vasoconstrictive effects in the coronary arteries and dose-dependently increases coronary perfusion pressure in a calcium-dependent manner thereby enhancing arteriole contractility^27^. The density of GHSR is increased in the atherosclerotic coronary artery and the saphenous vein compared to the corresponding non-diseased artery and vein^28^, indicating that up-regulation of ghrelin-GHSR signalling may occur within the diseased vasculature. In the same study, no differences in GHSR density were observed in the LA or the LV between the ischemic heart disease or dilated cardiomyopathy to the control tissues^28^. Our findings also showed no changes in myocardial GHSR as measured by Cy5-ghrelin(1-19), and a relatively weak correlation with ghrelin, suggesting that more dramatic changes in GHSR may occur at the site of vascular tissue injury or dysfunction.

The secretion of both natriuretic peptide type-B (BNP) and its N-terminal form (NT-proBNP), the current “gold standard” clinical heart failure (HF) biomarkers, are increased under conditions of cardiomyocyte stress and pressure overload. Circulating BNP levels are also increased in patients with aortic stenosis and mitral regurgitation, and can be used as a potential indication for valve replacement in patients with normal ejection fraction^29^. However, myocardial BNP mRNA does not change in patients who had aortic valve stenosis^30^. In our current study in patients with valvular disease, we also did not find a change in the tissue expression of BNP in either the LA or the LV. If tissue levels of BNP do not change, but levels of circulating BNP increase in valvular disease, it may suggest that the rate of post-translational processing of pro-BNP, rather than transcription or translation, increases in cardiomyocytes.

Interestingly, we found that BNP and ghrelin correlated in the left ventricle, and furthermore, strongly co-localized to the same intracellular compartment within ventricular cardiomyocytes. In healthy individuals, BNP is predominantly localized in the atrium; in heart failure, BNP appears to localize primarily in the ventricles^31^. In aortic stenosis, the endocrine profile of aortic valves, which includes natriuretic peptides, their receptors and processing enzymes, is altered ^30^; taken together with our results, we suggest that a regional shift in myocardial endocrine programming may be part of the progression of heart disease to heart failure. This regional shift may extend to other genetic programs as well; in dogs exposed to prolonged rapid ventricular pacing, there were drastic changes in genes expressing apoptosis, cell structure and mobility, and inflammation in the left atrium, while genes involved in metabolism and Ca^2+^ signalling changed only in the left ventricle ^32,33^. Therefore, there are clear regional differences in the genetic programs that underlie cardiomyocyte function in heart disease. Further studies will determine the intracellular dynamics of ghrelin and BNP during the progression of heart disease.

We also evaluated GHSR signalling pathways involving contractility and inflammation. We were particularly interested in correlations with SERCA2a, a common marker of contractility, as it has been shown to be activated through the GHSR signalling pathway, reducing intracellular Ca^2+^ levels and improving LVEF after myocardial infarction^2^. Additionally, we have previously found strong relationships between ghrelin and SERCA2a in both a mouse model of cardiomyopathy^13^ and human heart failure^11^. In the present study, we observed a positive correlation between SERCA2a and GHSR-ghrelin only in the left ventricle, which may indicate the importance of contractility signalling in the left ventricle, as it is the predominant chamber for contractile force. Additionally, SERCA2a and BNP were also correlated only in the left ventricle. Through binding natriuretic peptide receptor-A, BNP activates cGMP signalling, which is coupled to L-type calcium channels and intracellular calcium levels through SERCA2a in ventricular cardiomyocytes^34^. As LVEF was unchanged in most of the patients, it is likely that alterations in the relationships between SERCA2a, ghrelin, and BNP within left ventricular cardiomyocytes occurs prior to any overt functional changes as detected by echocardiography.

Toll-like receptor 4 (TLR4) is a marker of inflammation elevated in the myocardium in end stage heart failure^35,36^, and ghrelin has been shown to attenuate the pro-inflammatory effects of TLR4 in HF^4,37^. In this study, we did not find any significant correlations between TLR4 and GHSR or ghrelin or other biomarkers, which may indicate the TLR4-mediated pro-inflammatory axis is not co-regulated with the ghrelin-GHSR system at this point in valvular heart disease.

Another measure of cardiac damage is the presence of fibrosis (collagen and fibroblast deposition) in the myocardium, which increases with HD severity^38,39^. Our results indicate a significant increase in the fibrotic deposition in only the left atrium of HD patients when compared to the control heart tissues. The apparent lack of left ventricular fibrosis is consistent with the preserved LVEF of this cohort^40^. This accumulation of fibrosis present in the left atrium may be associated to the downstream signalling pathway changes seen in GHSR-ghrelin in this cohort as increased fibrosis deposition is common in the heart during the later stages of heart failure and is a sign of cardiomyocyte damage. However, these results are a bit difficult to interpret due to the large degree of variability in fibrosis in the patient cohort. This result is not surprising due to the heterogenous nature of fibrosis deposition in the myocardium and the subsequent difficulty in obtaining representative biopsy specimens^41^.

There are a few limitations to our study to note. Our study was conducted with a small sample size that limited effective disaggregation of the data by region. It is important to note that the control and diseased tissues were embedded differently (paraffin wax vs fixed frozen respectively), although all samples were stained with the same protocols for each marker tested. Quantification of both paraffin and frozen healthy heart tissue is shown in supplemental figure S2 where there was no difference in the quantification between either tissue storage method. In addition, control LV tissue was obtained up to 1-3 days post-mortem which could allow for potential protein degradation between time of death and when the tissue samples were put in fixative during autopsy. Finally, we did not obtain blood samples from the patients, which would have given useful insight into the relationship between myocardial and circulating levels of ghrelin and BNP.

To conclude, we have identified changes in the myocardial GHSR-ghrelin axis in human valvular heart disease as compared to control cardiac tissues. The correlation between ghrelin and BNP and the colocalization in the myocardium of these peptides may suggest a regional shift in myocardial endocrine programming of cardiac cells in valvular heart disease. We have shown that the contractility marker SERCA2a was selectively correlated in the left ventricle, potentially as a cardioprotective mechanism prior to decreased LVEF. Our ongoing work will help to characterize GHSR-associated biochemical changes associated with early stages of heart disease.

## 5. Acknowledgements

We would like to thank Dr. and Peter Plugfelder and Ms. Anna Mcdonald for assistance in the collection of endomyocardial biopsies, and the Pathology Laboratory Core Department at the London Health Sciences Center for autopsy samples and their assistance in fibrosis staining of all tissue samples.

---

Supplemental Figure S1. Punctate staining with RenyiEntropy algorithm. Top panel shows representative fluorescent staining of healthy, diseased left atrium (LA) and diseased left ventricle (LV) tissue. Bottom panels show what the RenyiEntrophy algorithm considers positive staining (green) and what is background staining (black) for all types of tissues.

Supplemental Figure S2. Comparison between paraffin embedded control human tissue and frozen healthy heart biopsy samples. Marker comparison between GHSR (A), Ghrelin (B), and BNP (C) indicated no significant difference in quantification between methods of tissue storage.

## References

1. Ma, Y., Zhang, L., Launikonis, B. S. & Chen, C. Growth hormone secretagogues preserve the electrophysiological properties of mouse cardiomyocytes isolated from in Vitro ischemia/reperfusion heart. Endocrinology 153, 5480–5490 (2012).

2. Ma, Y., Zhang, L., Edwards, J. N., Launikonis, B. S. & Chen, C. Growth hormone secretagogues protect mouse cardiomyocytes from in vitro ischemia/reperfusion injury through regulation of intracellular calcium. PLoS One 7, (2012).

3. Raghay, K., Akki, R., Bensaid, D. & Errami, M. Ghrelin as an anti-inflammatory and protective agent in Ischemia/Reperfusion injury. Peptides 124, (2019).

4. Wang, Q. et al. Ghrelin protects the heart against ischemia/reperfusion injury via inhibition of TLR4/NLRP3 inflammasome pathway. Life Sci. 186, 50–58 (2017).

5. Baldanzi, G. et al. Ghrelin and des-acyl ghrelin inhibit cell death in cardiomyocytes and endothelial cells through ERK1/2 and PI 3-kinase/AKT. J. Cell Biol. 159, 1029–1037 (2002).

6. Huang, C.-X. et al. Ghrelin inhibits post-infarct myocardial remodeling and improves cardiac function through anti-inflammation effect. Peptides 30, 2286–2291 (2009).

7. Yang, C., Liu, Z., Liu, K. & Yang, P. Mechanisms of ghrelin anti-heart failure: Inhibition of Ang II-induced cardiomyocyte apoptosis by down-regulating AT1R expression. PLoS One 9, (2014).

8. Wang, Q. et al. Ghrelin Ameliorates Angiotensin II-Induced Myocardial Fibrosis by Upregulating Peroxisome Proliferator-Activated Receptor Gamma in Young Male Rats. Biomed Res. Int. 2018, 1–14 (2018).

9. Yang, C. et al. Ghrelin suppresses cardiac fibrosis of post-myocardial infarction heart failure rats by adjusting the activin A-follistatin imbalance. Peptides 99, 27–35 (2018).

10. Beiras-Fernandez, A. et al. Altered myocardial expression of ghrelin and its receptor (GHSR-1a) in patients with severe heart failure. Peptides 31, 2222–8 (2010).

11. Sullivan, R. et al. Dynamics of the Ghrelin/Growth Hormone Secretagogue Receptor System in the Human Heart Before and After Cardiac Transplantation. J. Endocr. Soc. 3, 748–762 (2019).

12. Aleksova et al. Ghrelin Derangements in Idiopathic Dilated Cardiomyopathy: Impact of Myocardial Disease Duration and Left Ventricular Ejection Fraction. J. Clin. Med. 8, 1152–1172 (2019).

13. Sullivan, R. et al. Changes in the Cardiac GHSR1a-Ghrelin System Correlate With Myocardial Dysfunction in Diabetic Cardiomyopathy in Mice. J. Endocr. Soc. 2, 178–189 (2018).

14. Pei, X. M. et al. Protective effects of desacyl ghrelin on diabetic cardiomyopathy. Acta Diabetol. 52, 293–306 (2015).

15. Fox, K. F. et al. Coronary artery disease as the cause of incident heart failure in the population. Eur. Heart J. 22, 228–236 (2001).

16. Kadoglou, N. P. E. et al. Serum levels of apelin and ghrelin in patients with acute coronary syndromes and established coronary artery disease-KOZANI STUDY. Transl. Res. 155, 238–246 (2010).

17. Zhang, M. et al. Plasma ghrelin levels are closely associated with severity and morphology of angiographically-detected coronary atherosclerosis in Chineses patients with diabetes mellitus. Acta Pharmacol. Sin. 33, 452–458 (2012).

18. Wang, F. et al. Ghrelin reduces rat myocardial calcification induced by nicotine and vitamin D3 in vivo. Int. J. Mol. Med. 28, 513–519 (2011).

19. Li, G. Z. et al. Ghrelin blunted vascular calcification in vivo and in vitro in rats. Regul. Pept. 129, 167–176 (2005).

20. McGirr, R., McFarland, M. S., McTavish, J., Luyt, L. G. & Dhanvantari, S. Design and characterization of a fluorescent ghrelin analog for imaging the growth hormone secretagogue receptor 1a. Regul Pept. 172, 69–76 (2011).

21. Douglas, G. A. F. et al. Characterization of a far-red analog of ghrelin for imaging GHS-R in P19-derived cardiomyocytes. Regul Pept. 54, 81–88 (2014).

22. Locatelli, V. et al. Growth hormone-independent cardioprotective effects of hexarelin in the rat. Endocrinology 140, 4024–4031 (1999).

23. Kennedy, D. J. et al. Central role for the cardiotonic steroid marinobufagenin in the pathogenesis of experimental uremic cardiomyopathy. Hypertension 47, 488–495 (2006).

24. Braunwald, E. et al. The Hemodynamics of the Left Side of the Heart as Studied by Simultaneous Left Atrial, Left Ventricular, and Aortic Pres-sures; Particular Reference to Mitral Stenosis. Circulation XII, 69–81 (1955).

25. Schwarz, F., Flameng, W., Schaper, J. & Hehrlein, F. Correlation between myocardial structure and diastolic properties of the heart in chronic aortic valve disease: Effects of corrective surgery. Am. J. Cardiol. 42, 895–903 (1978).

26. Papotti, M. et al. Growth hormone secretagogue binding sites in peripheral human tissues. J. Clin. Endocrinol. Metab. 85, 3803–3807 (2000).

27. Pemberton, C. J. et al. Ghrelin induces vasoconstriction in the rat coronary vasculature without altering cardiac peptide secretion. Am. J. Physiol. - Hear. Circ. Physiol. 287, 1522–1529 (2004).

28. Katugampola, S. D., Pallikaros, Z. & Davenport, A. P. [125I-His9]-Ghrelin, a novel radioligand for localizing GHS orphan receptors in human and rat tissue; up-regulation of receptors with atherosclerosis. Br. J. Pharmacol. 134, 143–149 (2001).

29. Bergler-Klein, J., Gyöngyösi, M. & Maurer, G. The Role of biomarkers in valvular heart disease: Focus on natriuretic peptides. Can. J. Cardiol. 30, 1027–1034 (2014).

30. Peltonen, T. O. et al. Distinct downregulation of C-type natriuretic peptide system in human aortic valve stenosis. Circulation 116, 1283–1289 (2007).

31. Hystad, M. E. et al. Regional cardiac expression and concentration of natriuretic peptides in patients with severe chronic heart failure. Acta Physiol. Scand. 171, 395–403 (2001).

32. Cardin, S. et al. Marked differences between atrial and ventricular gene-expression remodeling in dogs with experimental heart failure. J. Mol. Cell. Cardiol. 45, 821–831 (2008).

33. Pelouch, V., Milerová, M., Ošťádal, B., Hučín, B. & Šamánek, M. Differences between atrial and ventricular protein profiling in children with congenital heart disease. Mol. Cell. Biochem. 147, 43–49 (1995).

34. Rose, R. A. & Giles, W. R. Natriuretic peptide C receptor signalling in the heart and vasculature. J. Physiol. 586, 353–366 (2008).

35. Yu, L. & Feng, Z. The Role of Toll-Like Receptor Signaling in the Progression of Heart Failure. Mediators Inflamm. 2018, 1–11 (2018).

36. Yang, Y. et al. The emerging role of toll-like receptor 4 in myocardial inflammation. Cell Death and Disease vol. 7 1–10 (2016).

37. Liu, S. P. et al. Octanoylated Ghrelin Inhibits the Activation of the Palmitic Acid-Induced TLR4/NF-κ B Signaling Pathway in THP-1 Macrophages. ISRN Endocrinol. 2012, 1–8 (2012).

38. Barasch, E. et al. The Association Between Elevated Fibrosis Markers and Heart Failure in the Elderly: The Cardiovascular Health Study. Circ. Hear. Fail. 2, 303–10 (2009).

39. Travers, J., Kamal, F., Robbins, J., Yutzey, K. & Blaxall, B. Cardiac fibrosis. Circ. Res. 118, 1021–1040 (2016).

40. Burlew, B. S. & Weber, K. T. Cardiac fibrosis as a cause of diastolic dysfunction. PLoS One 27, 92–98 (2002).

41. Nagaraju, C. K. et al. Myofibroblast Phenotype and Reversibility of Fibrosis in Patients With End-Stage Heart Failure. J. Am. Coll. Cardiol. 73, 2267–2282 (2019).

