## Supplementary figures and images for "Regional Differences in the Ghrelin-Growth Hormone Secretagogue Receptor Signalling System in Human Heart Disease"

### Supplemental figures 1 and 2

Supplemental Figure S1.

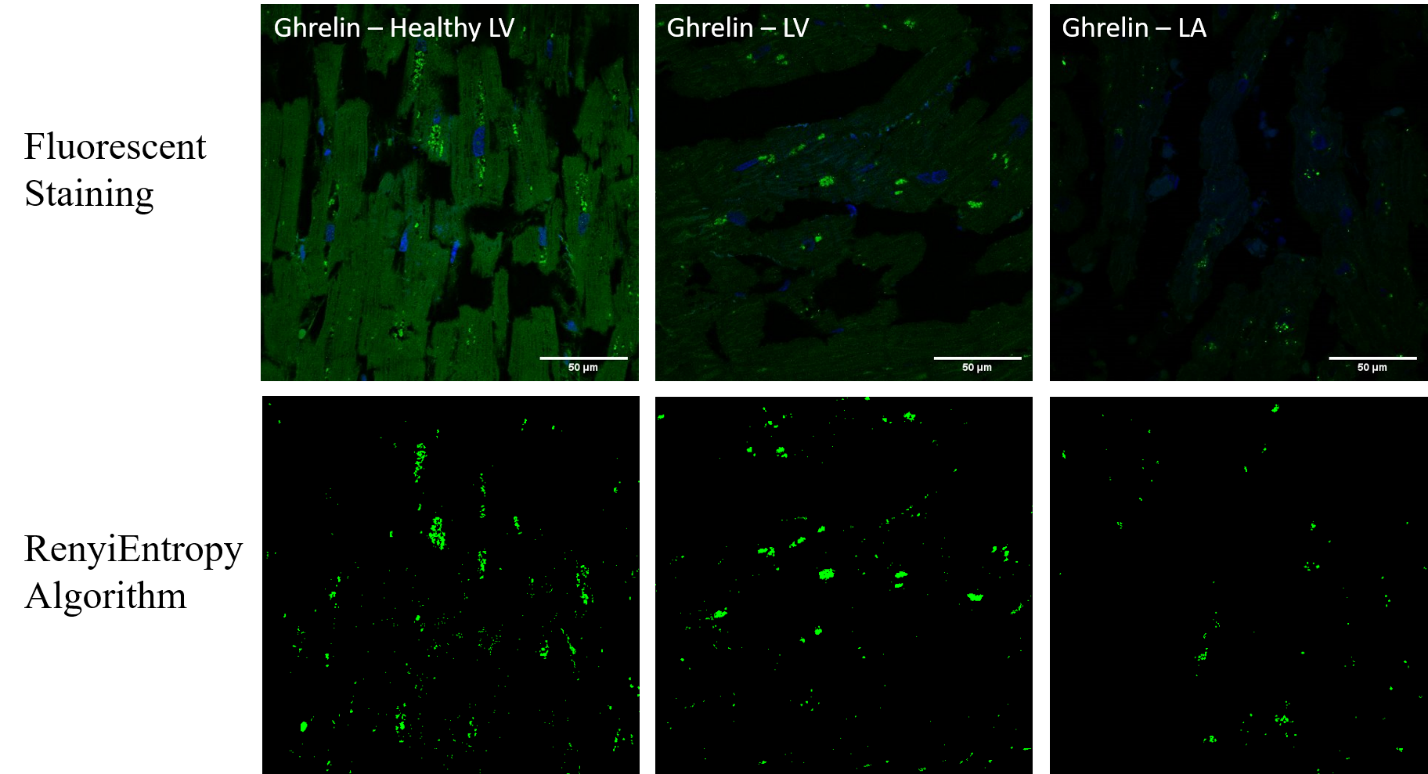

Supplemental Figure S2.

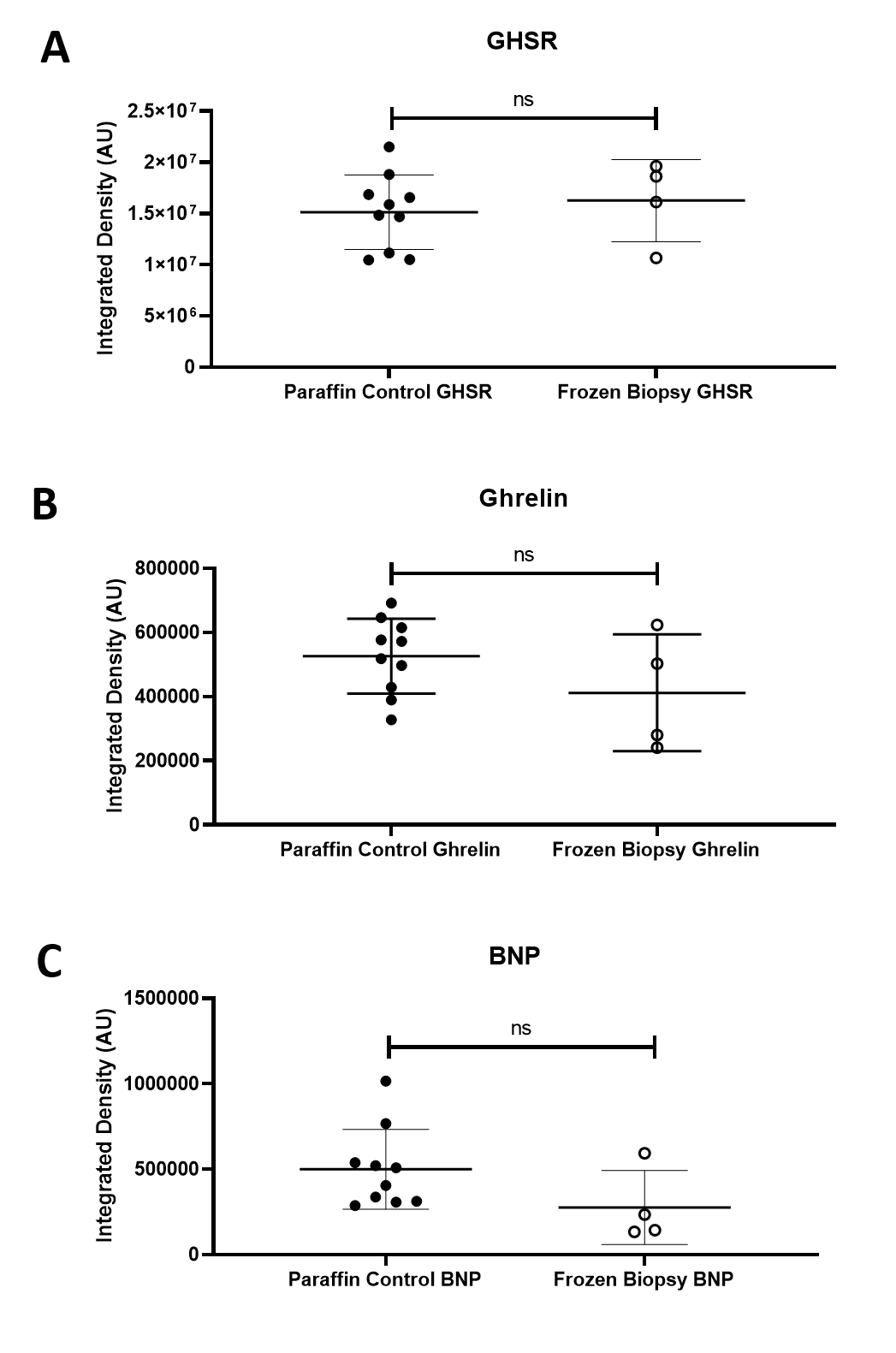
